# The adaptive architecture is shaped by population ancestry and not by selection regime

**DOI:** 10.1101/2020.06.25.170878

**Authors:** Kathrin A. Otte, Viola Nolte, François Mallard, Christian Schlötterer

## Abstract

Understanding the genetic architecture of adaptive phenotypes is a key question in evolutionary biology. One particularly promising approach is Evolve and Resequence (E&R), which combines advantages of experimental evolution such as time series, replicate populations and controlled environmental conditions, with whole genome sequencing. The recent analysis of replicate populations from two different *Drosophila simulans* founder populations, which were adapting to the same novel hot environment, uncovered very different architectures - either many selection targets with large heterogeneity among replicates or fewer selection targets with a consistent response among replicates. Here, we exposed the founder population from Portugal to a cold temperature regime. Although almost no selection targets were shared between the hot and cold selection regime, the adaptive architecture was similar: we identified a moderate number of loci under strong selection (19 selected alleles, mean selection coefficient = 0.072) and very parallel responses in the cold evolved replicates. This similarity across different environments indicates that the adaptive architecture depends more on the ancestry of the founder population than the specific selection regime. These observations have a pronounced impact on our understanding of adaptation in natural populations.

## Introduction

Adaptation of natural populations to environmental change may either occur from standing genetic variation or by the acquisition of new mutations. The relative importance of these two paths crucially depends on the underlying adaptive architecture (Barghi et al. 2020) of the focal trait. The adaptive architecture differs from the genetic architecture, which is inferred by QTL mapping and GWAS, by accounting for pleiotropic constraints as well as for the large body of deleterious mutations (Gazal et al. 2017; Zeng et al. 2018). Depending on the context, two different aspects of the adaptive architecture are emphasized. The focus is either the identity of specific loci/SNPs or the general characteristics of the adaptive architecture characterized by the number of contributing loci and their effect sizes and frequency in the focal population (Barghi et al. 2020).

Contributing loci are either identified by QTL/GWAS studies (Flint and Mott 2001; McCarthy et al. 2008) or with genomic selection scans, which apply statistical tests to detect selection signatures from population polymorphism data (Storz 2005; Vitti et al. 2013). Some selection scans assume that selection targets are shared among populations exposed to the same environment, because such parallel signatures provide additional statistical support (Turner et al. 2010; Lee and Coop 2017; Harris and DeGiorgio 2020). Many selection targets were successfully identified with these strategies and have contributed to our understanding of the molecular basis of adaptation processes (e.g. Turner et al. 2010; Jones et al. 2012; Roesti et al. 2014; Reid et al. 2016; Johnson and Voight 2018). It is, nevertheless, not apparent to what extent these results can be generalized, because most adaptive traits have a polygenic basis (Barton and Keightley 2002; Sella and Barton 2019) and either only small allele frequency changes (Sella and Barton 2019) or non-parallel responses are expected (Barghi et al. 2020).

The key concept of polygenic adaptation is that multiple loci are contributing to the phenotype, and rather than focusing on particular loci, the collective effect of all loci needs to be considered to estimate the phenotypic value of a given trait. This has important implications for the understanding of the adaptive architecture (Barghi et al. 2020).

The infinitesimal model (Fisher 1918; Bulmer 1971; Barton et al. 2017) is the most extreme case of polygenic adaptation and is frequently approximated by very many contributing loci, each of very small effect. When many loci are contributing to a phenotype under stabilizing selection, any selection regime changing the trait optimum will result only in very small allele frequency shifts (Bulmer 1971; Sella and Barton 2019) - almost impossible to detect with classic population genetic tests (Pritchard et al. 2010; Field et al. 2016; Jain and Stephan 2017a).

Even when these conditions are relaxed and a distribution of effect sizes with some large effect alleles is considered, no pronounced allele frequency changes are expected when the populations are large and in mutation selection equilibrium: alleles with large effects are segregating at low frequencies only and do not contribute much to the phenotypic variance of the population upon which selection is operating (de Vladar and Barton 2014; Jain and Stephan 2017b). Theory predicts that as the pool of contributing loci to the selected phenotype becomes smaller (i.e. a decreased mutational target), larger allele frequency changes are expected that will progressively be detected in population genetic analyses (Höllinger et al. 2019). Therefore, traits with an intermediate number of contributing loci are particularly interesting, because the response of these loci can be sufficiently strong to be detected in experiments while, at the same time, being informative about polygenic adaptation: more loci are segregating in the population than required to reach a new trait optimum (i.e. genetic redundancy).

The consequence of this genetic redundancy is that the contribution to the phenotype can be highly heterogeneous for individual loci in differentiated populations if they vary in frequency. This expectation nicely conforms with empirical data, mostly from QTL studies, which find heterogeneous sets of contributing loci among different populations (Adeyemo et al. 2009; Wu et al. 2013; Al Olama et al. 2014; Li and Keating 2014; Conte et al. 2015; Horikoshi et al. 2018; Takata et al. 2019; Wojcik et al. 2019; Zan and Carlborg 2019; Hodonsky et al. 2020). In the case of adaptation to a new trait optimum, genetically differentiated populations will adapt by frequency changes of different sets of loci. Hence, for polygenic adaptation the identity of individual selected loci is not very important to describe the adaptive architecture, rather information about the number of loci, effect sizes and frequencies are needed to understand the selective response.

Selection signatures not shared among natural populations are difficult to interpret, as the distinction between population-specific selection targets and false positive/negative signals can be challenging given the high impact of a largely obscure demography on selection signatures (Jensen et al. 2005; Stajich and Hahn 2005; Li et al. 2012; Lohmueller 2014; Pavlidis and Alachiotis 2017; Johri et al. 2020). Experimental evolution, in contrast, provides the advantage of replicate populations, which evolve from the same founder population under controlled experimental conditions (Kawecki et al. 2012). The potential of experimental evolution to study the genomic signatures of polygenic adaptation has, however, not yet been fully exploited since most studies apply truncating selection. Thus, the contributing alleles experience continued selection pressure throughout the entire experiment, causing a parallel selection response in the replicate populations towards an extreme phenotype. Laboratory natural selection is a specific experimental evolution design, where the evolving populations are exposed to a new environment (Garland and Rose 2009). In contrast to truncating selection, populations are expected to reach a new phenotypic optimum. In combination with whole genome sequencing, it provides an interesting approach to study the adaptive architecture experimentally.

Two previous experimental evolution studies conducted in the same novel hot laboratory environment revealed very different adaptive architectures (Mallard et al. 2018; Barghi et al. 2019). In the Portugal experiment, five strongly selected genomic regions were identified and this selection signal was highly parallel across replicates. For the Florida experiment, 99 selection targets were identified and considerable heterogeneity was observed between the replicates. One possible explanation for this different adaptive architecture is that the ancestral trait optima differed between the two founder populations (Barghi and Schlötterer 2020), leading to a more intense selection in the Portugal experiment, because it was less well adapted than the Florida founder population to high temperatures. Alternatively, more large effect alleles may have been segregating at higher frequencies in the Portugal founder population.

Here, we exposed replicate populations of the Portugal founders to a cold temperature regime to shed more light on the different selection responses. Interestingly, we found very little overlap between the genomic position of the selection targets in the hot and cold temperature regimes. Most large effect loci detected in the hot environment did not respond in the cold, suggesting that hot and cold temperature adaptation may be different traits, rather than a simple shift in optimum of the trait ‘temperature adaptation’. Nevertheless, adaptation to both, hot and cold, temperature regimes had a very similar adaptive architecture - with a comparable number of selection targets and effect sizes. We conclude that the adaptive architecture differs between populations and may be trait independent. We discuss to what extent this phenomenon can be explained by the infinitesimal model.

## Results

We studied the genetic architecture of cold adaptation in *Drosophila simulans* by combining experimental evolution with whole genome re-sequencing (Evolve and Resequence, E&R). Five replicate populations originating from the Portuguese founder population described by Mallard et al. (2018) evolved for more than 50 generations (about four years) in a cold temperature regime with daily fluctuations between 10°C and 20°C. Genome-wide allele frequencies were determined in 10 generation intervals by sequencing pools of individuals (Pool-Seq (Schlötterer et al. 2014)). Contrasting generation 0 with 51 we identified 6,527 SNPs, which changed in frequency more than expected by genetic drift either across all five replicates (adapted CMH test (Spitzer et al. 2020), 6,510 SNPs) or at least in one replicate (adapted χ^2^ test (Spitzer et al. 2020), additional 17 SNPs). The X chromosome harbored only 142 SNPs. Such a low number of candidate SNPs on this chromosome was not seen in other *Drosophila* E&R studies that observed similar numbers of candidate SNPs on the X chromosome and autosomes (Jha et al. 2015; Jha et al. 2016; Barghi et al. 2019; Kelly and Hughes 2019; Michalak et al. 2019). The pronounced peak structure in the Manhattan plot (Figure 1A) indicates that many candidate SNPs are not independent due to linkage (Nuzhdin and Turner 2013; Franssen et al. 2017a). We accounted for this and employed a correlation-based haplotype reconstruction approach to identify independently selected haplotype blocks based on their distinct trajectories (Franssen et al. 2017a; Otte and Schlötterer 2017) and treated each of these haplotype blocks as a single target of selection (Barghi et al. 2019).

**Figure 1:**
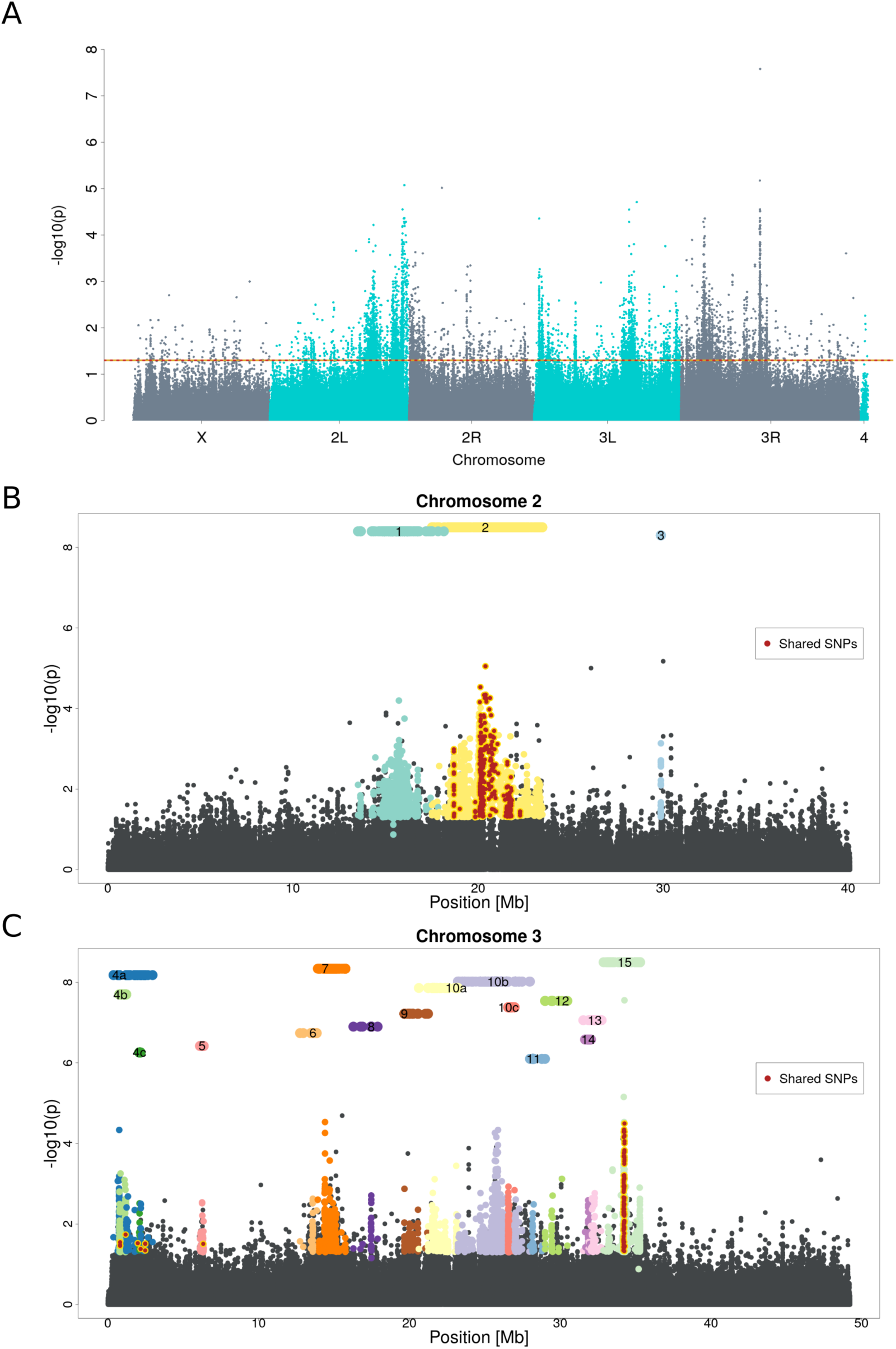
Manhattan plots of the genomic selection signature in response to cold temperature. A) p-values were obtained from an adapted CMH test (Spitzer et al. 2020) comparing the founder generation (F0) to the most advanced (F51) generation. The dotted line indicates the significance threshold (p-value < 0.05 after correction for multiple testing). B, C) A close up of the Manhattan plot in panel A for chromosomes 2 and 3. Each selected haplotype block and the corresponding candidate SNPs are shown in a distinct color. Numbered bars above the Manhattan plot indicate the position of the selected haplotype blocks. Some blocks, which can be further subdivided by the analysis of earlier time points, are labeled with letters (e.g. 10a, 10b etc.). SNPs with significant allele frequency changes in both, hot and cold, selection regimes (shared SNPs) are colored in dark red.

### Selected haplotype blocks in the cold-evolved *D. simulans* population

Because the small number of candidate SNPs precluded haplotype block reconstruction on the X chromosome, all 15 haplotype blocks were identified on the major autosomes (Figure 1B, 1C). The size of the haplotype blocks ranged from 5 kb to 6 Mb. Three of the 15 blocks were located on chromosome 2 while all others fell on chromosome 3.

The 15 selected haplotype blocks were identified by clustering SNPs with similar allele frequency trajectories in the five replicates and six time points. A conservative assumption is that each of the selected haplotype blocks contains one selection target. Nevertheless, multiple selection targets can recombine during the experiment onto a single haplotype block, which outcompetes the individual parental haplotype blocks (Otte and Schlötterer 2019). Such multiple-target haplotypes will dominate the later phases of the experiment and are considered as a single selected haplotype. To identify such cases, we repeated the haplotype block reconstruction with fewer generations - four time points up to generation 31 - and found that two reconstructed haplotype blocks could be further broken up: block 4 and 10 were split into three different blocks each (4a, 4b, 4c and 10a, 10b, 10c, see Figure 1C). This observation confirms that the number of inferred independent haplotype blocks is a conservative estimate of the number of selection targets, and we used the sub-blocks for subsequent analyses whenever applicable, analyzing a total of 19 selected alleles.

Selected haplotype blocks are characterized by a set of marker SNPs which show correlated allele frequency trajectories across replicates. Nevertheless, the correlation is not very stringent to account for sequence diversity among the haplotypes carrying the selection target (Otte and Schlötterer 2019). Hence, not all of the haplotype block marker SNPs describe the frequency trajectory of the (unidentified) selection target equally well. Reasoning that SNPs with the most pronounced allele frequency change are the best representatives of the selection target, we used the 10% most significant marker SNPs of each haplotype block and refer to them as the selected allele. The frequency of each selected allele at every time point is determined as the median frequency of these 10% most significant SNPs. We used only replicates with a selection coefficient large enough to be significantly different from neutrality (p-value < 0.05), therefore we excluded replicates in which a given selected allele did not increase in frequency. The starting frequencies were highly variable among the 19 selected alleles. We detected selected alleles with starting frequency as low as 0.06, but also as high as 0.4 (Figure 2A). The selection coefficients were rather high and ranged from 0.04 to 0.11 (Figure 2A).

**Figure 2:**
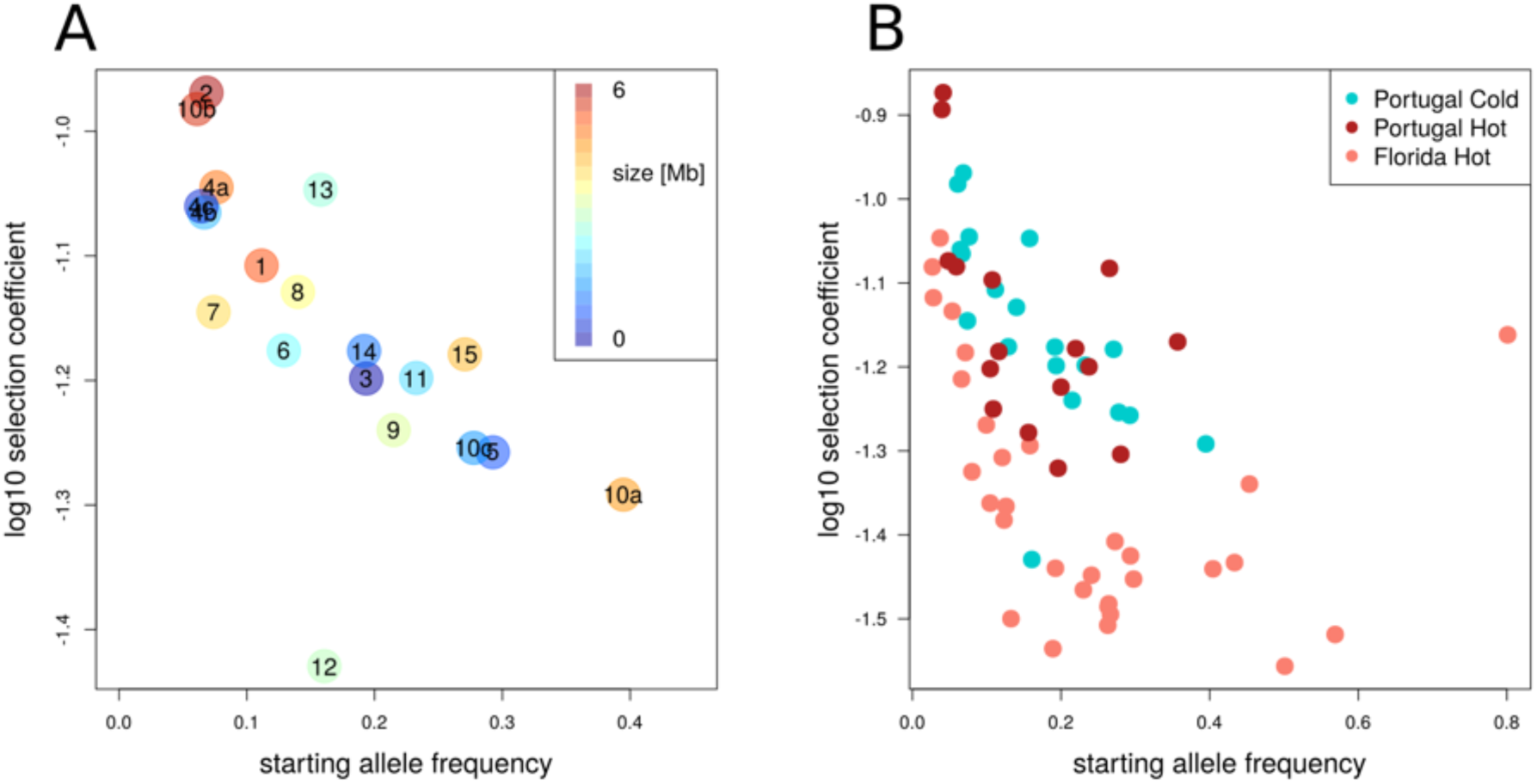
Inverse relationship between starting allele frequency and selection strength. For all experiments, the selected alleles were identified using the same protocol and are based on the top 10% SNPs in reconstructed haplotypes. A) The relationship is shown for the Portugal cold experiment. The color code reflects the size of the selected haplotype block, and starting allele frequencies and selection coefficient of the selection targets are plotted on x- and y-axis. The numbers relate to the selected haplotype blocks from Figure 1B and 1C with numbers indicating blocks detected in the analysis using all time points. Sub-blocks resulting from an analysis using earlier time points are labeled with the letters a, b or c. B) A qualitatively similar relationship between starting allele frequency of each haplotype block and selection strength is found in the cold-evolved Portugal (blue), hot-evolved Portugal (red) and hot-evolved Florida (pink) population. Nevertheless, the distribution in the Florida experiment was shifted towards lower selection coefficients while the two temperature regimes in the Portugal population were highly similar. We conclude that the adaptive architecture is population-specific but does not depend on the temperature regime.

Overall, we noticed a striking relationship between starting frequency and selection coefficients. Selected alleles starting with lower frequencies had higher selection coefficients than selected alleles with higher starting frequencies. This relationship was significant when analyzing the full set of 19 blocks including the broken-up haplotype blocks (i.e. replacing block 4 and 10 by sub-blocks 4a, 4b, 4c and 10a, 10b, 10c) and was not significantly influenced by block size (linear regression; factor: starting allele frequency p = 0.004 and factor: block size p = 0.130). The inverse relationship between starting allele frequency and selection coefficient was robust with respect to the definition of a selected allele (see Supplementary Figure S1).

While a negative correlation between frequency and effect size is expected by theory and has been previously reported for GWAS (Eyre-Walker 2010; Simons et al. 2014; Mancuso et al. 2016; Zeng et al. 2018) and E&R (Barghi et al. 2019) studies, it is important to note that a Beavis-like effect (Beavis 1998; Xu 2003) may also contribute to this observation: alleles with low starting frequencies require stronger selection to result in detectable allele frequency changes than alleles starting from intermediate frequencies.

With a median size of 1.5 Mb, the selected haplotype blocks were rather large. The median number of genes per selected haplotype block is 20 but it can reach up to 154 (see Supplementary Figure S2) in the largest reconstructed block (block 2, see Figure 1B). The smallest block contained only a single candidate gene (block number 3, 4.8 kb). All 23 marker SNPs were located within one intronic region of the gene *M-spondin* (*mspo*, FBgn0020269), an extra-cellular matrix protein of *Drosophila*, putatively involved in muscle development (Bataille et al. 2010). The role of this gene in temperature adaption is not apparent and further studies are required for a better understanding of this selection signature.

### Temperature-specific adaptation

Replicates from the same Portuguese founder population were also exposed to a hot selection regime fluctuating between 18°C and 28°C (Mallard et al. 2018). Both temperature regimes have the same daily temperature amplitude of 10°C (cold 10/20°C and hot 18/28°C), but mean temperatures differ (15°C in the cold and 23°C in the hot regime). The temperatures were chosen such that one of the temperatures is stressful, whereas the other temperature is benign (David 1983; Petavy et al. 2001). Contrasting the founder population with hot-evolved generation 59, Mallard et al. (2018) identified few (five) very pronounced selection peaks, some of them related to metabolic alterations in the hot-evolved populations.

For an unbiased comparison of the two experiments, we added time series data for the hot-evolved populations (F0, F15, F37, F59) and applied the same haplotype reconstruction pipeline as described above. Similar to the cold-evolved population, the X chromosome had too few outlier SNPs (114 SNPs) for haplotype reconstruction. 16 selected haplotype blocks were identified on the two major autosomes (Supplementary Figure S3) and their selection coefficients ranged from 0.05 to 0.13 (Figure 2B). It is remarkable that not only the number of inferred selection targets, but also the distribution of selection coefficients is highly similar for the two temperature regimes (hot-evolved = 16 blocks, cold-evolved = 19 blocks).

Only two haplotype blocks shared more SNPs than expected by chance between the two temperature regimes. Furthermore, the shared region was in both cases only a small part of the total haplotype block (Figure 1B and 1C, blocks 2 and 15). A prominent similarity between the selection regimes could be identified in block 15, where the majority of overlapping SNPs were located in the gene *Ace* (FBgn000024). Nevertheless, the selection pattern for *Ace* differs between both temperature regimes (Langmüller et al. 2020). The shared SNPs of block 2 were located in several genes, and therefore no clear candidate for common adaptation could be identified in this region.

We further scrutinized the haplotype blocks that were not shared between the selection regimes and had a starting frequency higher than 0.15 to rule out that a selection signature in opposite direction - as expected for a polygenic trait selected in contrasting environments - was missed. The allele frequency change of all candidate SNPs in a haplotype block was always higher in the focal temperature regime. Importantly, in both, hot and cold, selection regimes we very rarely observed a frequency change in the opposite direction (Supplementary Figure S4). We conclude, therefore, that we have no support for alleles being selected in opposite direction in hot and cold temperatures. Rather, most alleles show a temperature-specific response.

### Population-specific adaptation

Independent of the temperature regime the evolved populations derived from the Portugal founder population revealed only a moderate number of selection targets. This contrasts a recent experiment using the same hot temperature regime but a founder population from Florida (Barghi et al. 2019). For a consistent comparison to the Portugal population, we repeated the analysis of the Florida population using our haplotype reconstruction pipeline but focused only on the two major autosomes. 31 selected haplotype blocks were identified on chromosome 2 and 3 (Supplementary Figure S5), which are considerably fewer selected alleles than the 88 reported by Barghi et al. (2019) for these two chromosomes. This difference reflects an alternative strategy to identify candidate SNP sets for the haplotype reconstruction rather than the clustering method (Otte and Schlötterer 2019). Following the same protocol as for the Portugal experiments, 9,197 outlier SNPs were identified, which is more conservative than the 52,199 outlier SNPs used by Barghi et al. (2019) for haplotype reconstruction. The Florida population harbored about twice as many selected haplotype blocks as the hot-evolved Portugal population. For all experiments we identified the same relationship between starting frequency and selection coefficients, but the distribution for Portugal was shifted towards higher selection coefficients (Figure 2B). This result is robust with respect to the definition of selected alleles (see Supplementary Figure S6). This difference in selection coefficients remains significant when we account for allele frequencies in the founder populations (contrasts between estimated marginal means Portugal cold - Florida hot p < 0.0001 and Portugal hot - Florida hot p < 0.0001). No significant difference was observed between the two temperature regimes of the Portugal population (Portugal cold - Portugal hot p=0.98).

The Florida experiment was based on twice the number of replicates as the Portugal experiment. To rule out that the number of replicates affects the inferred adaptive architecture, we repeated the analysis with 100 sets of five randomly sampled Florida replicates. In all 100 random subsets the selection coefficient was not different from the full data set (Wilcoxon rank-sum test p > 0.05 adjusted for multiple testing using the Benjamini-Hochberg method), suggesting that the difference between the Portugal and Florida data sets cannot be explained by a different number of replicates.

We quantified the degree of parallelism between the experiments using the Jaccard index on the selected haplotype blocks with *s* significantly different from zero. High values indicate parallel genetic responses whereas low values reflect heterogeneous, non-parallel genetic responses between replicates. Jaccard indices were high for the two Portugal experiments (median similarity between replicates 80% and 82%, respectively). For the Florida experiment, the Jaccard index was significantly lower (median similarity between replicates 70% for the full data set; Wilcoxon rank-sum test p-value < 0.001 against each Portugal data set), indicating less parallel genetic responses and therefore increased realized genetic redundancy. This pattern was robust with respect to the method used to define a selected allele in a given replicate (Figure 3 and Supplementary Figure S7).

**Figure 3:**
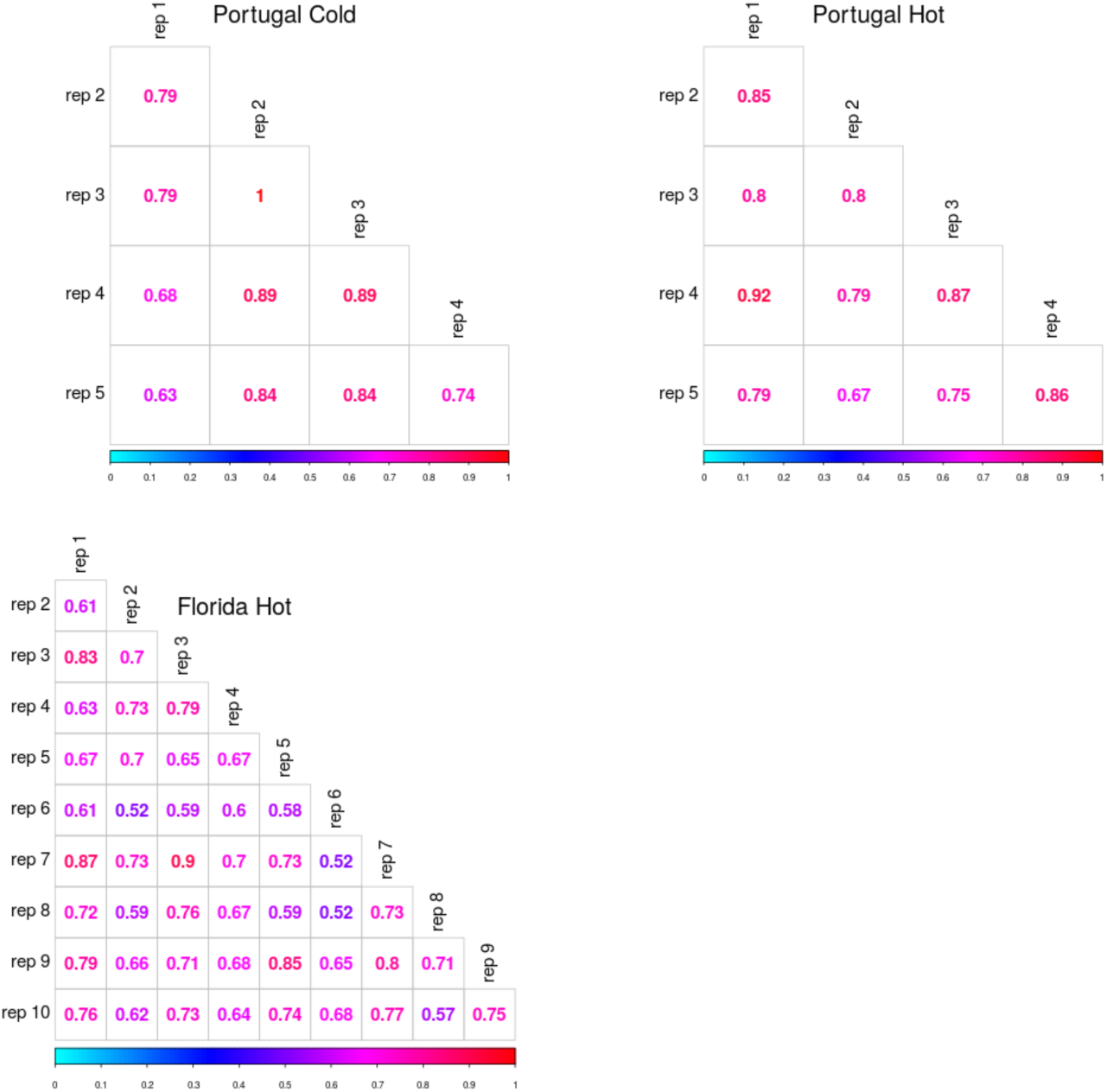
More parallel selection signatures among evolved replicate populations derived from Portugal founders than from those with Florida ancestry. Jaccard indices comparing the different replicates in the cold-evolved Portugal (top left), hot-evolved Portugal (top right) and hot-evolved Florida (bottom) population. Jaccard index was computed based on estimation of significant selection coefficients (p-value < 0.05).

The comparison of selected haplotype blocks between evolved populations derived from different founder populations is difficult because the selected haplotype blocks are reconstructed only for significant SNPs. Even if the selected haplotype block is shared, the remaining haplotypes differ between the populations. As the frequency change at a given SNP on the selected haplotype depends on the frequency of this SNP in the non-selected haplotypes, different candidate SNPs are being identified - even if the same haplotype block is selected.

Thus, the same selected haplotype block may have different marker SNPs in two different populations - suggesting that different haplotype blocks are selected. Furthermore, the low linkage disequilibrium in natural *Drosophila* populations (Charlesworth and Charlesworth 1973; Langley et al. 1974) implies that very few haplotypes are expected to be shared between samples from different populations. For these reasons, we did not attempt to test whether the same haplotype blocks are selected in Portugal and Florida.

## Discussion

*D. simulans* populations of different origin (Portugal and Florida) had very distinct adaptive architectures in the same hot temperature regime (Mallard et al. 2018; Barghi et al. 2019): Portugal had fewer selection targets, which were strongly selected in a highly parallel manner. Florida harbored more selection targets with more heterogeneity among replicates and lower selection coefficients compared to Portugal. In this report, we studied replicate populations derived from the Portugal founder population, which adapted to a cold temperature regime to understand why such different adaptive architectures were inferred in these two populations. Below, we discuss several possible explanations for the differences in adaptive architecture.

### Different trait optima in the ancestral populations

Both founder populations were collected on different locations with their own specific temperature profile and in different phases of the seasonal cycle (Portugal in July 2008, Florida in November 2010). Assuming that temperature adaptation is a single high-level trait, the ancestral trait optimum may differ on the phenotypic axis. This implies that a population, which is less adapted to hot environments should be better adapted to cold environments. Less well adapted populations will experience stronger and more parallel selection responses across replicates (Franssen et al. 2017b; Barghi et al. 2020), consistent with the pattern observed in the hot Portugal population. Different trait optima of the founder populations were further supported by the observation that the Portugal founder population is less fecund than the Florida founder population when assayed in the novel hot environment (Barghi et al., unpublished results). This implies that the mean phenotype of the Portugal population is more distant from the new trait optimum in the hot laboratory environment than the Florida population.

The analysis of the cold-evolved replicates casts some doubts on this simple interpretation. We assumed that the hot and cold experiments shifted the trait optimum into opposite directions relative to the (unknown) trait optimum of the ancestral Portugal population. Hence, contributing alleles segregating at sufficiently high frequency in the ancestral population should be selected in opposite direction in the two temperature regimes. Nevertheless, the results did not fit our expectations - most selected haplotype blocks were not shared between the two temperature regimes. While selected haplotype blocks starting from low frequencies may not be identified as selection targets in the opposite temperature regime, also haplotype blocks with higher allele frequencies in the founder population were not selected in opposite direction.

Hence, we conclude that temperature adaptation may not be a single high-level phenotype. Rather, several sub-phenotypes on a lower level, which are not all shared for the different temperature regimes, are contributing to adaptation. This conclusion is further supported by different genomic signatures of hot and cold stressors in E&R (Tobler et al. 2014) and QTL mapping (Morgan and Mackay 2006) studies.

### Differences in adaptive variation

Autosomal polymorphism levels differ between the two founder populations with Florida being more variable than Portugal (π_Florida_ = 0.0076 and π_Portugal_ = 0.0062, Wilcoxon rank-sum test on non-overlapping 10 kb windows, p-value < 0.001). Assuming that neutral variability is a good approximation of adaptive variation, which is not always the case (Kellermann et al. 2009), Portugal is expected to harbor less adaptive variation than Florida. This implies that Florida reaches the trait optimum faster than Portugal (Thornton 2019; Barghi and Schlötterer 2020), but in absence of phenotypic time series data, we cannot assess this hypothesis. A particularly interesting hypothesis related to the different polymorphism levels is that Portugal harbors so little adaptive variation that it does not have much genetic redundancy. This would imply that no (or only limited) excess of adaptive genetic variation is segregating in the Portugal founder population that can be used to reach the trait optimum.

The Florida founder population, in contrast, harbors a considerable excess. Such differences in the number of contributing loci can generate quite different patterns of parallel selection responses (Barghi and Schlötterer 2020), matching the Portugal and Florida experiments. Nevertheless, it is not apparent that the moderate differences in genome-wide polymorphism levels are sufficiently large to explain this pattern.

### Linkage disequilibrium

The above discussion about the heterogeneity of the inferred genetic architectures between populations and selection regimes rests on the central assumption that the major contributing loci were identified and could be distinguished with a recently developed haplotype reconstruction approach (Otte and Schlötterer 2019). In other words, it is assumed that only a moderate number of distinct loci contribute to adaptation.

Alternatively, the observed selection response may be explained by many loci of small effect - an idea that matches in its extreme form the infinitesimal model (Barton et al., 2017). Empirical support for a highly polygenic architecture of many traits comes from the strong correlation between chromosome length and the fraction of heritability explained (Visscher et al. 2007; Yang et al. 2011; Shi et al. 2016). If multiple small effect loci cluster together this may result in a signature that will be interpreted as a single selection target (Yeaman and Whitlock 2011). Short genomic segments with a local clustering of favored loci can even introgress and leave the strong selection signature of a local allele frequency change (Sachdeva and Barton 2018). Empirical support for the clustering of contributing loci comes from the molecular dissection of candidate loci identified in QTL mapping studies. Single QTL loci can be broken into multiple SNPs contributing to the corresponding trait (Stam and Laurie 1996; King et al. 2012; Kerdaffrec et al. 2016; Gibert et al. 2017; Zan et al. 2017; Shahandeh and Turner 2020).

The situation in polymorphic founder populations is significantly more complicated than the simple two genotype case studied by Sachdeva and Barton (2018), but we propose that blocks of linked loci can not only generate pronounced selection signatures, but may also explain the differences in adaptive architecture between the Portugal and Florida experiment. Depending on the extent of linkage disequilibrium (LD) the clustering of contributing loci can vary. Hence, populations with different levels of LD may also harbor more or less clustered contributing loci. The influence of haplotype structure can be illustrated by two extreme cases: in the case of complete linkage equilibrium (LE), in any genomic window the haplotypes segregating in the population should have similar fitness despite being highly diverse. As a consequence, changes in trait optimum will result only in rather small frequency changes of the haplotypes in this genomic window. This pattern becomes more pronounced with an increasing number of contributing loci.

On the other hand, in the presence of strong linkage disequilibrium, fewer distinct haplotypes are present in a given genomic window. Sampling variation in the ancestral population generates haplotypes with different numbers of contributing loci in a given genomic window. The more pronounced the difference in the number of loci among haplotypes in a genomic window is, the stronger will be the fitness differences among them and thus the allele frequency change after a shift in trait optimum. Hence, because the difference in the number of contributing loci among haplotypes differs among genomic windows, linkage disequilibrium generates heterogeneity in selection response along the chromosome.

We illustrated the impact of LD by assuming about 1200 contributing loci genome-wide and simulated a window size of 1 Mb in a typical E&R setting. For high LD and LE the same number of chromosomes with beneficial alleles was used. 100 different genomic windows were simulated and simulations with higher LD resulted in a more heterogeneous response to selection. Consistent with a larger phenotypic variance (Figure 4A) also much more pronounced allele frequency changes were observed for some windows in the presence of linkage disequilibrium (Figure 4B). Hence, while very homogeneous moderate frequency shifts were observed for beneficial alleles in linkage equilibrium, some pronounced sweep windows were detected for windows with LD. We also assessed the degree of parallelism in the response between replicates and found a more parallel response for the high LD simulations (Figure 4C).

**Figure 4:**
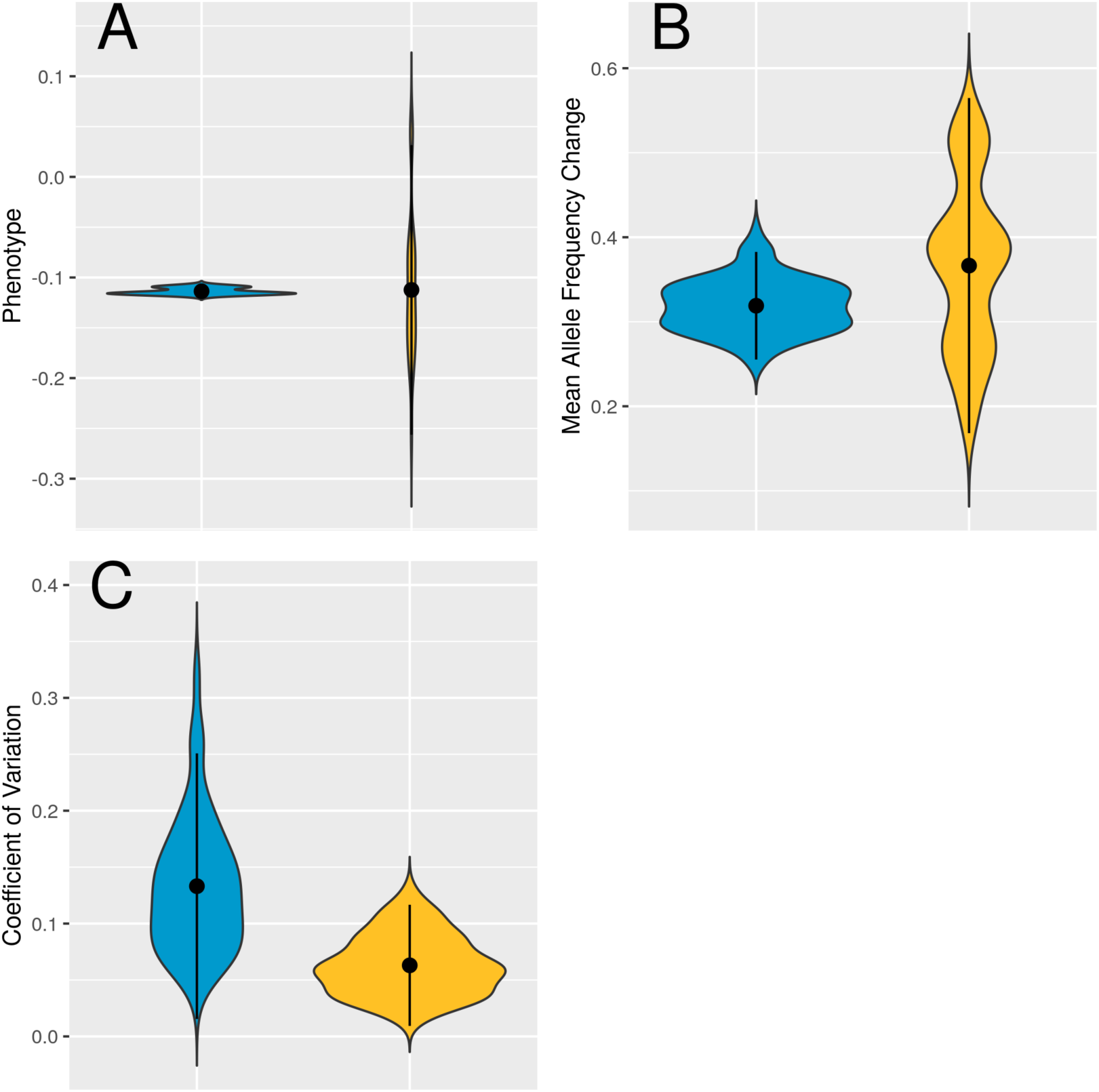
Influence of linkage disequilibrium on the genomic response of a polygenic trait. We simulated 50 generations of polygenic adaptation after a shift in trait optimum. 100 replicates, each with 10 loci in a 1 Mb region either with linkage equilibrium (blue) or strong LD (yellow) are shown in violin plots. A) Independent of linkage structure, the same mean phenotype was reached, but simulations with high LD were considerably more scattered. B) Pronounced allele frequency changes were obtained for both, linkage equilibrium und high LD. For linkage equilibrium, the genomic windows were all rather similar, indicating that no window showed a strong selection signature distinguishing it from the genomic background. Simulations with strong LD, however, resulted in highly heterogeneous selection responses, with some windows having a frequency change larger than 0.6, which is a strong selection signature distinguishing it from the remaining genomic windows. C) The heterogeneity among five replicate populations is measured by the coefficient of variation. The selection response in simulations with linkage equilibrium was less parallel than in those with strong LD.

Thus, for polygenic traits the inferred adaptive architecture can be strongly affected by linkage in the ancestral population. A previous study on the impact of recombination on the response to a shift in trait optimum with truncating selection observed more heterogeneity among replicates in the case of linkage equilibrium than for complete linkage (Zhang and Hill 2005). We attribute these differences to the small population sizes in Zhang and Hill (2005).

Nevertheless, does this scenario of a highly polygenic architecture with differences in LD apply to the Florida and Portugal experiments? Following the same rationale as Shi et al. (2016), Barghi et al. (2019) tested whether longer haplotype blocks were more strongly selected than shorter ones, but no significant correlation was found. Similar results were observed for the cold evolved Portugal populations. We caution, however, that these negative results do not provide strong support for the identification of distinct selection targets. Possible, not mutually exclusive, reasons for the lack of significance even in the presence of a highly polygenic architecture are: 1) the contributing loci have different effect sizes, thus small haplotype blocks with large effect loci may be more strongly selected ones than larger blocks, 2) larger blocks may harbor more loci with effects in opposite direction than smaller ones, 3) since haplotype blocks are still relatively short (compared to full chromosomes) stochastic sampling and heterogeneity in the density of contributing loci may obscure the correlation between size of the selected haplotype block and the selection response.

The hypothesis that LD differences can explain the heterogeneity in inferred adaptive architecture is supported by the observation that the two founder populations differed across all chromosomes in their pattern of linkage disequilibrium (Figure 5). The higher LD in Portugal compared to Florida is fully consistent with the prediction that founder populations with higher LD are more likely to show stronger and more parallel sweep signatures than populations with lower LD. Although this pattern fits our observations, in absence of more information about the degree of polygenicity and the distribution of contributing loci across the chromosomes and their effect sizes, it is not possible to determine whether the observed differences in LD are sufficient to explain our empirical data.

**Figure 5:**
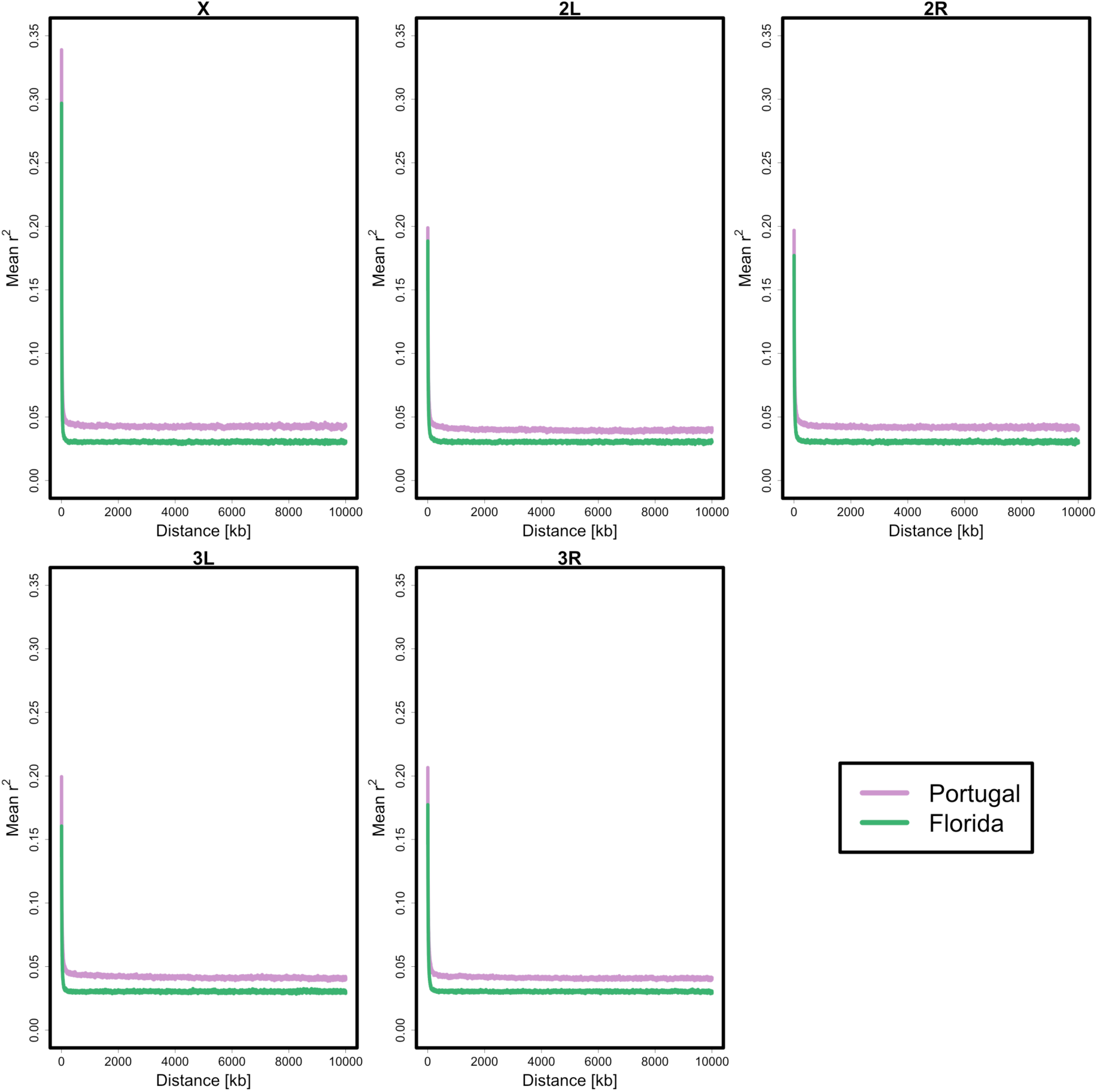
Linkage disequilibrium in the ancestral Portugal und Florida population as measured by the mean r^2^ of loci with distances up to 10,000 kb based on 34 individual haplotype sequences.

### Temperature adaptation may involve multiple, temperature-specific traits

While we cannot pinpoint the cause for the differences in the inferred adaptive architecture between Portugal and Florida, the analysis of cold-evolved replicates shed some important light on our understanding of temperature adaptation.

Many studies, in particular theoretical ones, considered high level phenotypes, such as temperature adaptation, as a single adaptive trait, where shifts in mean temperature are treated as a simple shift in trait optimum (e. g. Bridle et al. 2009; Chevin et al. 2010; Hoffmann 2010; Kopp and Matuszewski 2014). This implies that all segregating contributing loci affect the optimal phenotype - irrespective of the position of the optimum - i.e. in the case of temperature adaptation irrespective of whether the optimum is in the hot or cold. The limited overlap between the selection targets in hot- and cold-evolved replicates is striking, as it contradicts this assumption. With the exception of the genomic region around the *Ace* locus and a region across the centromere of chromosome 2, which changed in the same direction in both temperature regimes, no shared haplotype blocks were detected. To some extent, the lack of shared haplotype blocks can be attributed to low starting frequencies, which implies that selection in the opposite direction does not result in allele frequency changes sufficiently large to be detected.

Nevertheless, even for haplotype blocks starting from intermediate frequency, no selection signature in the opposite direction was noticed.

This implies that different loci are contributing to adaptation in hot and cold environments - irrespective of whether a highly or moderately polygenic architecture is assumed. A very similar lack of shared candidates was also noticed in a *D. melanogaster* experiment, where replicate populations were exposed to the same hot and cold temperature regimes (Tobler et al. 2014). This experiment was, however, conducted for a much smaller number of generations, and the selection signature was analyzed on the SNP-level, which makes the interpretation of the results particularly challenging given the contribution of large segregating inversions to temperature adaptation in this species (Hoffmann et al. 2002; Rako et al. 2007).

The observation of different selection targets in hot and cold environments is particularly interesting, because seasonal changes were found to be associated with cycling allele frequencies in natural *D. melanogaster* populations (Bergland et al. 2014), which suggests that the same SNPs are being selected in opposite direction in hot and cold environments. One possible explanation for these differences to our study is that in natural populations only a moderate number of generations separates the two temperature regimes, while in our experiment the temperature regime remained constant across more than 50 generations.

### Population-specific adaptive architectures

Various studies, mainly using QTL mapping and GWAS, have identified different loci contributing to the same trait in diverged populations (Adeyemo et al. 2009; Wu et al. 2013; Al Olama et al. 2014; Li and Keating 2014; Conte et al. 2015; Horikoshi et al. 2018; Takata et al. 2019; Wojcik et al. 2019; Zan and Carlborg 2019; Hodonsky et al. 2020; Kemppainen et al. 2020). A recent experimental evolution study using *D. subobscura* populations with different genetic background also observed very little overlap in the genomic regions responding to a common selection regime (Seabra et al. 2018). Hence, the different selection targets obtained from Portugal and Florida experiments conducted in the same hot environment are not particularly surprising and emphasize the limited insights about the genetic basis of a polygenic trait from single population studies. Very surprising, however, was the observation that the adaptive architecture (number of contributing loci, effect sizes and starting frequencies) was different between Portugal and Florida, but strikingly similar between the hot and cold selection regime.

Although more experiments are needed to nail down why the adaptive architecture is highly dependent on the founder population and not on the selection regime, our results have important implications for all studies attempting to characterize the adaptive architecture. The analysis of a single population cannot be sufficient to understand the genetic basis of adaptive traits. Thus, multiple diverged populations need to be studied to reach conclusions that can be generalized beyond a limited number of focal populations.

## Material and Methods

Unless stated otherwise, analysis was conducted using R v3.6.1 (R Core Team 2019).

### Experimental Populations and Selection Regime

The set-up of the evolution experiment is described in detail elsewhere (Mallard et al. 2018). In brief, female flies were sampled from a natural *Drosophila simulans* population in Northern Portugal in the summer of 2008 and used to establish 250 isofemale lines. These lines were kept for 10 generations in the laboratory before starting the experiment. Mated females of all lines were used to create the starting populations. Ten replicates were created by combining an equal number of flies from each line. Five of the replicates were then kept in a 12h:12h day and night cycle with temperatures of 20°C during the day and 10°C during the night (*cold regime*). The population size was kept constant at 1250 per replicate in non-overlapping generations.

The other five replicates were used to start the evolution experiment described by Mallard et al. (2018), which was identical to the cold regime except for the temperature which fluctuated between 28°C during the day and 18°C at night (*hot regime*).

### Evolve & Resequence

For the founder population, we used the sequences described by Mallard et al. (2018), but added new sequence data from two replicates of generation F3 from the cold regime to increase coverage. To avoid biases related to different sequencing approaches, all reads were pooled and then randomly split into five subsets with a coverage of 100x each. These subsets were used as founder population replicates throughout the analysis of the cold-evolved and the (re-)analysis of the hot-evolved populations. While sequence data for the F59 in the hot regime were available from Mallard et al. (2018), we added new Pool-Seq data for the intermediate time points F15 and F37 in the hot regime to allow for the time series analysis and haplotype reconstruction in this study. All sequences used for the founder population (including the F3 from the cold regime) and all sequences of the hot regime were derived from females only.

Sequencing data from the time points F11, F21, F31, F41 and F51 in the cold regime were newly generated from pooled females and males. Paired-end libraries were generated with different protocols and sequenced on different Illumina platforms (see Supplementary Table S1).

### Data processing

Sequencing reads were trimmed using Readtools TrimFastq version 0.2. (Gomez-Sanchez and Schlötterer 2017) with the parameters --minReadLength 50 --disable5pTrim -- mottQualityThreshold 20. The *D. simulans* genome sequence created by Palmieri et al. (2015) was used as reference for read mapping. To avoid false positive outlier SNPs, which might arise when libraries with different read lengths and insert sizes are combined in one analysis (Kofler et al. 2016), three different mappers were used to map reads of the two time points used for outlier testing (cold regime: F0 and F51; hot regime: F0 and F59). Reads were mapped using Bowtie2 version 2.2.6 (Langmead and Salzberg 2012) with parameters --end-to-end --X 1500, bwa mem version 0.7.13 (Li and Durbin 2010) with default parameters and novoalign version 3.03.2 (Novocraft 2014) with parameters -i 350,100 -F STDFQ -o SAM -r RANDOM.

The intermediate time points (cold regime: F11, F21, F31, F41; hot regime F15, F37) used for the detection of selected haplotype blocks were mapped with novoalign only, as this mapper is known to estimate allele frequencies most accurately (Kofler et al. 2016).

Mapped reads were filtered for mapping quality ≥ 20 and proper pairs using SAMtools view version 1.3.1 (Li et al. 2009). Duplicates were removed using picard MarkDuplicates version 2.1.1 (Broad Institute 2019). Barcoded files were split using Readtools AssignReadGroupByBarcode version 0.2.2 (Gomez-Sanchez and Schlötterer 2017) with parameters --maximumMismatches 1. The BAM files were then used to create mpileup files with SAMtools mpileup version 1.3.1 (Li et al. 2009), and finally, PoPoolation2 mpileup2sync.jar version 1.201 (Kofler et al. 2011) was used to create sync files from mpileup files. All subsequent analysis was conducted on the basis of these sync files.

### SNP calling and masking

Single nucleotide polymorphisms (SNPs) were called from the founder population by creating sync files from BAM files as described above but filtering for polymorphic sites that had a mapping quality of at least 30 and a minimal count of at least 5 and were detected by the three mapping algorithms in all founder replicates. Filtering resulted in 3.8 million SNPs that were used for further analysis. Indels were detected using the PoPoolation2 identify-indel-regions.pl script, and transposable elements were detected with repeatmasker v 1.332 (Smit et al. 2015). Custom databases made by combining RepeatMasker database Dfam_Consensus-20181026, RepBase-20181026, and transposon_sequence_set.embl.txt from flybase.org (FB 2018_06), and search engine NCBI/RMBLAST v 2.2.27+ were used for repeats > 500 bp. All sync files were masked for these repetitive regions and for known Y chromosome translocations (Tobler et al. 2017) using the PoPoolation2 filter-sync-by-gtf.pl script.

### Correcting for different insert sizes

To correct for false positive outlier SNPs created by libraries with different insert sizes (Kofler et al. 2016), mapping results from the three different mappers for the founder (F0) and most evolved (cold regime: F51; hot regime F59) population were used. χ^2^ tests were conducted to compare the results of the different mappers per replicate and time point. After correcting for multiple testing using the Benjamini-Hochberg procedure (Benjamini and Hochberg 1995), only SNPs that showed a consistent response across comparisons (p.adjust ≥ 0.05) were kept for further analysis.

### Candidate SNPs

Candidate outlier SNPs were detected in the filtered sync files created from novoalign mapping results after correcting for false positive outliers as described above. Allele frequency changes between the founder and the most evolved population were analyzed using CMH and χ^2^ tests which are adapted for genetic drift and pool sequencing noise as implemented in the R package ACER version 1.0 (Spitzer et al. 2020). SNPs within the top 1% of coverage were excluded from the analysis to avoid copy number variants. Intermediate generations were included in the correction approach. Effective population size per replicate was calculated using the R package poolSeq version 0.3.5 (Taus et al. 2017) with the function estimateWndNe (window size *10 kb*, method *P.planI*, pool size and census size *1250*) and used for CMH and χ^2^ tests. The CMH test was performed using all population replicates per time point whereas χ^2^ tests were performed for each replicate separately. All results were corrected for multiple testing (Benjamini-Hochberg). Finally, candidate SNPs detected by either test (p.adjust < 0.05) were combined to include consistent responses across replicates (CMH test) and replicate-specific responses (χ^2^ test).

### Selected haplotype blocks

Selected haplotype blocks were reconstructed from candidate SNP allele frequency data of all time points and replicates using the R package haplovalidate with MNCS of 0.01 (Otte and Schlötterer 2019). Haplotype blocks were reconstructed for all time points. As haplotype blocks might contain more than one selected allele, early time points (cold regime: F11, F21 and F31; hot regime: F15 and F37) were used for fine-mapping of selected haplotype blocks (Otte and Schlötterer 2019). Here, the analysis detected the characteristic signal of reconstructed haplotype blocks with multiple selection targets, which is the presence of a single haplotype block in the most evolved generation but several reconstructed haplotype blocks when analyzing the early generations separately. Haplotype blocks from the early generations showing this pattern were included in the final analysis.

Selection coefficients for the detected haplotype blocks per replicate were computed using the allele frequency trajectories of the top 10% outlier SNPs based on CMH and χ^2^ test result and using the poolSeq v0.3.5 function estimateSH (method *LLS*) (Taus et al. 2017). Only selection coefficients with p-value < 0.05 were used to calculate the median selection coefficient for each selected allele. Relationship of selection coefficient, starting allele frequency and block size were tested using a linear model with log10 transformation of selection coefficients. To test the robustness of our definition for selected alleles, we repeated the estimation of selection coefficients using a) the top 20% SNPs or b) all SNPs that had an allele frequency change > 0.1 (Supplementary Figure S1). Candidate genes per block were detected from the gene annotation of the reference genome (Palmieri et al. 2015) including also SNPs 200 bp up- and downstream of the focal gene.

### Comparison to other experimentally evolved *D. simulans* populations

We used data from the same *D. simulans* population evolving under a hot temperature regime (Mallard et al. 2018) to contrast adaptation to different temperatures. For this population, we included two additional time points, so that four time points in total were available: F0, F15, F37, F59. The data set was filtered, candidate SNPs were detected and haplotypes were reconstructed in the same way as described above including the combined analysis of all (F0-F59) and early (F0-F37) time points. To estimate how many shared marker SNPs were expected by chance, we randomly sampled the number of shared SNPs from the haplotype blocks, calculated the fraction per haplotype block (N=10,000) and finally applied a 95 % cut-off.

In addition, a different published hot-evolved *D. simulans* population from Florida (Barghi et al. 2019) was used for the comparative analysis. The data set was filtered and candidate SNPs were detected in the same way as described above. As intermediate time points were available for this data set (every 10th generation from F0 to F60), haplovalidate (Otte and Schlötterer, 2019) with MNCS of 0.01 was used to detect selected haplotype blocks including the combined analysis of all (F0-F60) and early (F0-F30) time points as described for the Portugal population. Selection coefficients were computed as described above.

We fitted a linear model with log10 transformed selection coefficients as response and main effects of population as fixed categorical effect with three levels (Florida hot, Portugal hot, Portugal cold) and a linear and quadratic covariate for starting allele frequency, to account for their non-linear relationship with the response, as explanatory variables. Residuals from this model were normally distributed and displayed variance homogeneity. The model with linear and quadratic covariate for starting allele frequency fit significantly better than a model with only a linear term. Contrasts between populations were compared based on estimated marginal means (R package *emmeans*).

We compared the similarity of replicates calculating the Jaccard indices for the Portugal and the Florida population using the R package *philentropy* (Drost 2018). We created binary data based on the replicate-specific selection response, i.e. whether or not a significant selection coefficient could be estimated by the poolSeq package (see above) for the corresponding replicate and allele (p-value <0.05). Following Barghi et al. (2019), we created binary data by applying a cut-off of 0.1 to the median allele frequency change of selected alleles per replicate. Jaccard indices between populations were compared using the two-sample Wilcoxon rank-sum test. To analyze the effect of the different number of replicates (10 in the Florida and five in the Portugal population) we repeated the analysis of selection coefficients and Jaccard indices on a downsampled set of Florida replicates. For this analysis, we took 100 random samples of five replicates from the Florida population data set and repeated the analysis for each of them as described above.

### Nucleotide diversity in the ancestral populations

Nucleotide diversity (π) of each autosome in the ancestral populations was calculated from the allele frequency data using the formula of Tajima (1989). For maintaining a comparable number of low-frequency alleles we subsampled the Florida data set to five replicates. As different sets of five Florida replicates resulted in very consistent π estimates (data not shown) we only used one set for the direct comparison to Portugal.

### Linkage disequilibrium in the ancestral populations

To quantify linkage disequilibrium, we used 189 haplotype sequences of the Florida founder population (Howie et al. 2019) and 34 haplotype sequences of the Portugal founder population which are described in Langmüller et al. (2020). For maintaining a comparable number of low-frequency alleles, we subsampled the Florida data set to 34 haplotypes. We calculated the mean r^2^ for loci (minor allele count = 3, minimum SNP quality = 50) within 10,000 kb distance. As different sets of 34 Florida haplotypes resulted in very consistent mean estimates (see Supplementary Figure S8) we only used one set for the direct comparison to Portugal.

### Linkage disequilibrium simulations

We illustrated the effect of linkage equilibrium (LE) or strong linkage disequilibrium (LD) on polygenic adaptation after a shift in trait optimum with computer simulations using MimicrEE2 v208 (Vlachos and Kofler 2018) in qff mode. We used parameters that matched the Portugal *D. simulans* E&R experiment with five replicates, each starting with the same 1000 homozygous individuals which evolved for 50 generations. For computational simplicity, we assumed that all 10 loci contributing to the phenotype, each with a starting allele frequency of 0.05 and effect size of 0.05, are restricted to a 1 Mb region. 100 independent simulations were performed to mimic 100 different genomic regions. We used a Gaussian fitness function as previously described (e.g. Barghi and Schlötterer 2020): minimum fitness 0.5, maximum fitness 4.5, standard deviation of the phenotype 1.2, heritability of 0.5. The mean fitness of the ancestral population was −0.44 and the new trait optimum was 0.5. We used the average recombination rate of *D. simulans* (Dsim_recombination_map_LOESS_100kb_1.txt, (Howie et al. 2019)). We generated two different sets of founder populations, one with strong LD and one with linkage equilibrium (LE). Both sets of founder populations contained 200 chromosomes with favored alleles and 800 chromosomes without. For strong LD, four different sets of selected haplotypes were generated, and the number of contributing loci was randomly distributed between the four sets of selected haplotypes. Hence, 50 haplotypes had the same number of contributing loci, but due to stochastic sampling the number of contributing loci differs among the four sets of 50 haplotypes. To generate starting populations in LE, we randomly distributed the selected alleles across 200 haplotypes until each of the alleles had a final frequency of 0.05.

After 50 generations, we generated “Pool-seq data” with 50x coverage and added sequencing noise by binomial sampling based on the allele frequencies. We recorded the phenotypic and mean frequency change across loci and replicates as well as the coefficient of variation in the mean allele frequency change across the five replicates as an indicator for the degree of parallel response.

## Data Availability

Sequence data were deposited at the European Nucleotide Archive (ENA) under the accession number XXX. Population sync files, all results and scripts were deposited on Dryad Digital Repository XXX.

## Acknowledgments

We thank Neda Barghi for sharing unpublished fecundity data and for providing information concerning the hot-evolved *D. simulans* data set. Anna Langmüller, Thomas Taus and Claire Burny shared code to remove SNPs with allele frequency estimates that are sensitive to the insert sizes of NGS sequencing libraries. The authors also thank the members of the Institute of Population Genetics for discussion and support on the project. Special thanks to Marlies Dolezal for in depth statistical advice. KAO was supported by a DFG Research Fellowship (OT 532/1-1). FM was supported by a Marie Sklodowska-Curie Individual Fellowship (H2020-MSCA-IF-661149). CS was supported by the European Research Council grant “ArchAdapt” and the Austrian Science Funds (FWF, P27630, P29133). Illumina sequencing for a subset of the data was performed at the VBCF NGS Unit (www.viennabiocenter.org/facilities).

