## Supplementary Material for "The adaptive architecture is shaped by population ancestry and not by selection regime"

This file includes:

Supplementary Figures S1 - S8

The following Supplementary Table is available separately (as .xlsx file):

Supplementary Table S1: Details of DNA extraction, Illumina library preparation and sequencing for all samples used in this study.

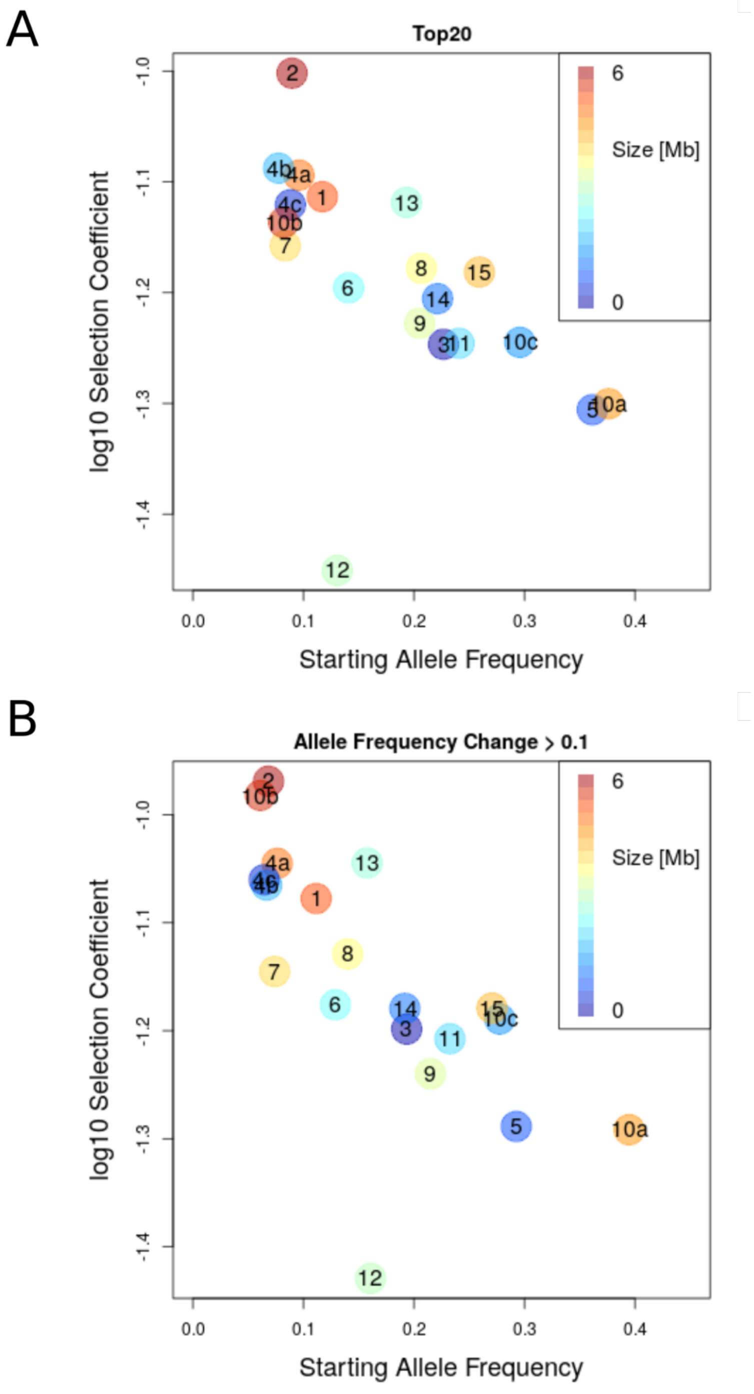


Supplementary Figure S1: The inverse relationship between starting allele frequency and selection coefficient is robust to to the definition of the selected allele. The color code reflects the size of the selected haplotype block, and starting allele frequencies and selection coefficient of the selection targets are plotted on x- and y-axis. The numbers refer to the selected haplotype blocks from Figure 1B and 1C in the main text. A) Selected allele defined by calculating selection coefficients from the top 20 % SNPs. B) Selected allele defined by calculating selection coefficients from all SNPs that have an allele frequency change > 0.1 estimated for the founder and the most evolved population.

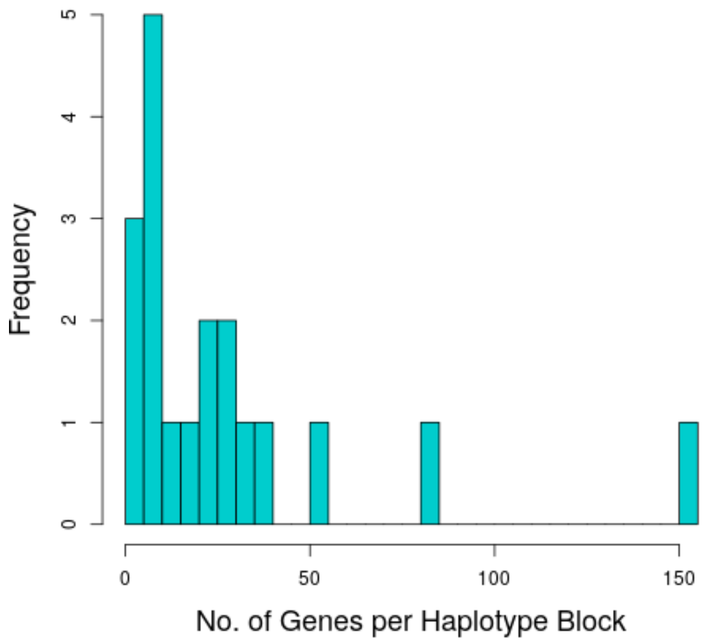


Supplementary Figure S2: Histogram of the number of genes per reconstructed haplotype block in the cold-evolved Portugal population.

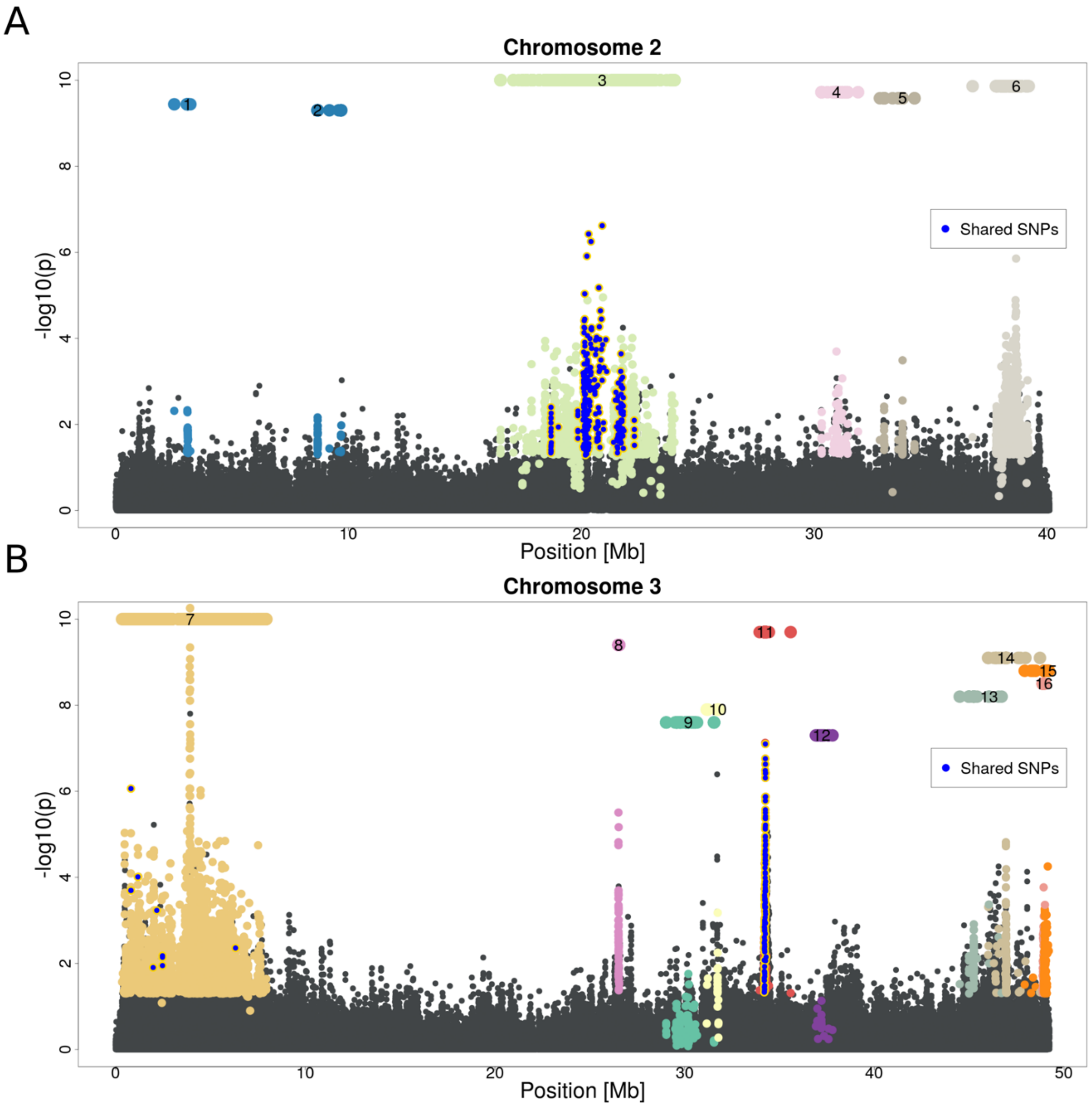


Supplementary Figure S3: Selected haplotype blocks from replicated *D. simulans* populations from Portugal evolving in a novel hot environment. A Manhattan plot of p-values obtained from an adapted CMH test for A) chromosome 2 and B) chromosome 3 is shown. Each selected haplotype block and the corresponding candidate SNPs are shown in a distinct color. Numbered bars above the Manhattan plot indicate the position of the selected haplotype blocks. SNPs with significant allele frequency changes in both, hot and cold, selection regimes (shared SNPs) are colored in blue.

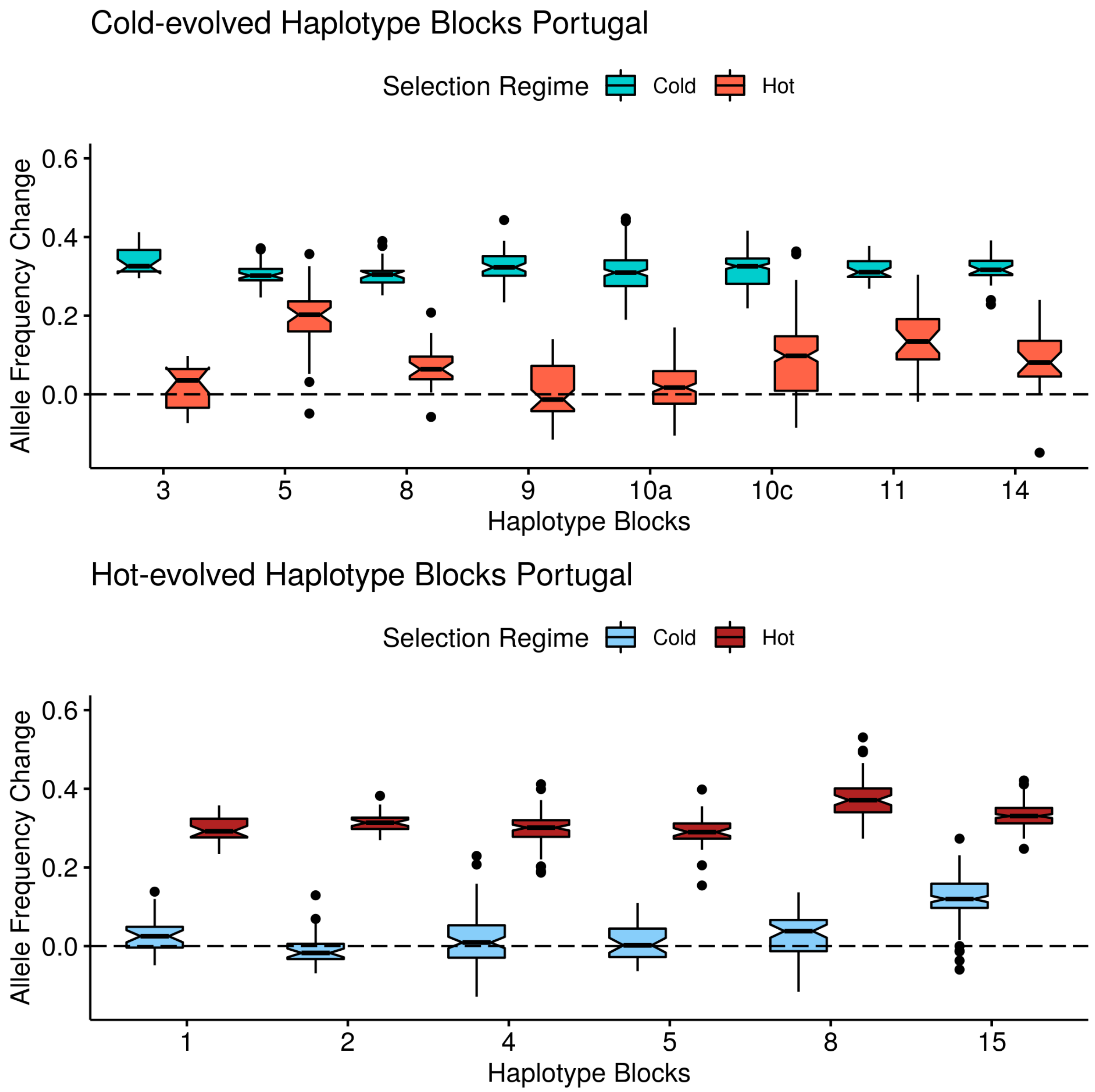


Supplementary Figure S4: Allele frequency changes between the founder and the evolved populations: all SNPs in the cold-evolved haplotype blocks (top) and hot-evolved haplotype blocks (bottom) of the Portugal experiment are shown. We only analyzed blocks that were not shared between the two selection regimes and had a starting allele frequency > 0.15. The upper panel shows candidate SNPs from selected haplotype blocks identified in the experiment with the cold temperature regime. While the candidate SNPs from the haplotype blocks selected in the cold (turquoise) show a strong frequency change, the same SNPs in the hot experiment (orange) showed no clear response or moved in the same direction as in the cold regime. The lower panel shows candidate SNPs in the haplotype blocks from the hot environment (red). Similar to the upper panel, the same SNPs in the cold experiment (blue) did not experience a pronounced frequency change in the opposite direction. Hence, this analysis does not support selection of the same allele in opposite direction in the hot and cold temperature regime, as expected for a single trait responding to a shift in trait optimum.

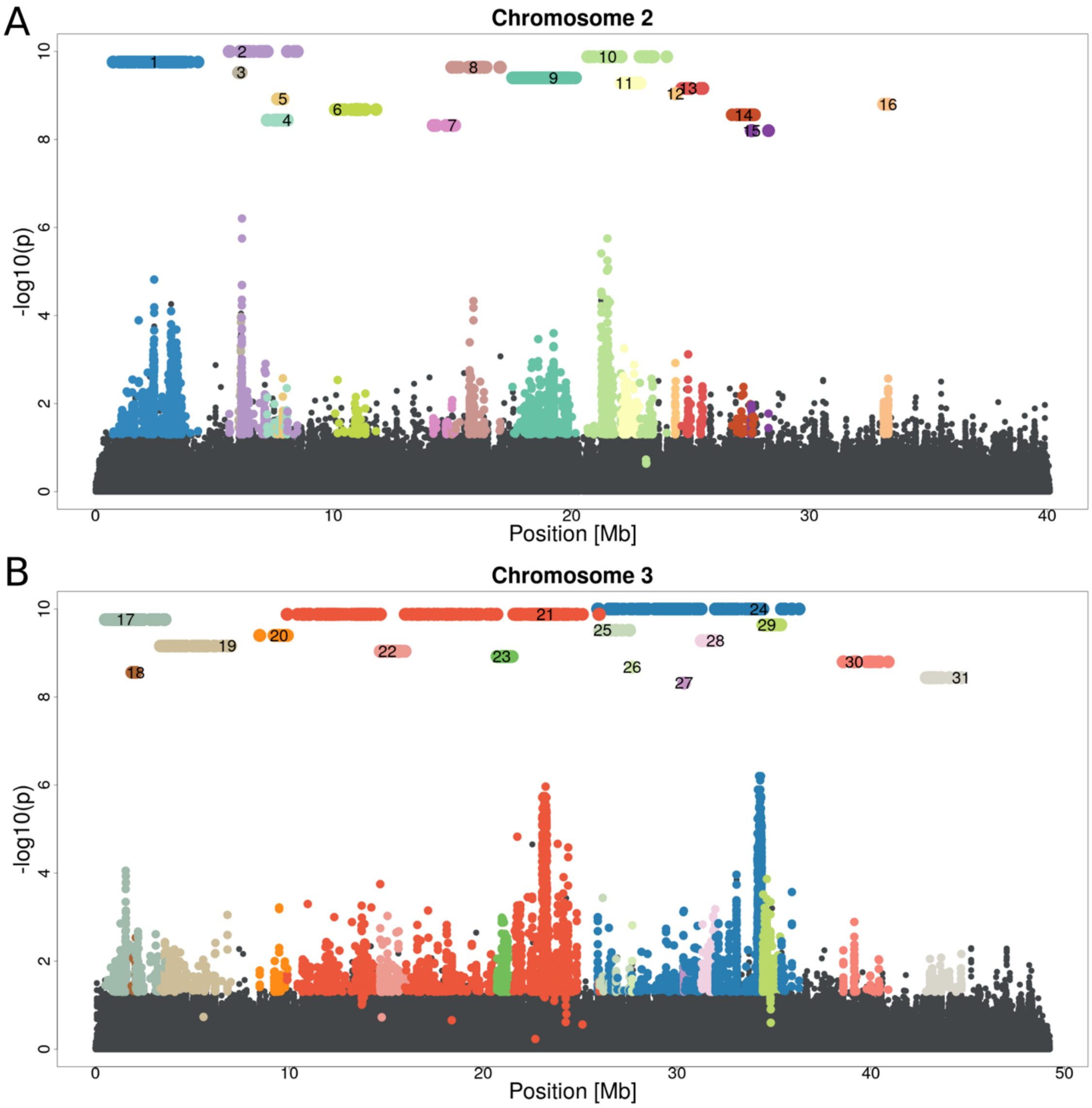


Supplementary Figure S5: Selected haplotype blocks from replicated *D. simulans* populations from Florida evolving in a novel hot environment. A Manhattan plot of p-values obtained from an adapted CMH test for A) chromosome 2 and B) chromosome 3 is shown. Each selected haplotype block and the corresponding candidate SNPs are shown in a distinct color. Numbered bars above the Manhattan plot indicate the position of the selected haplotype blocks.

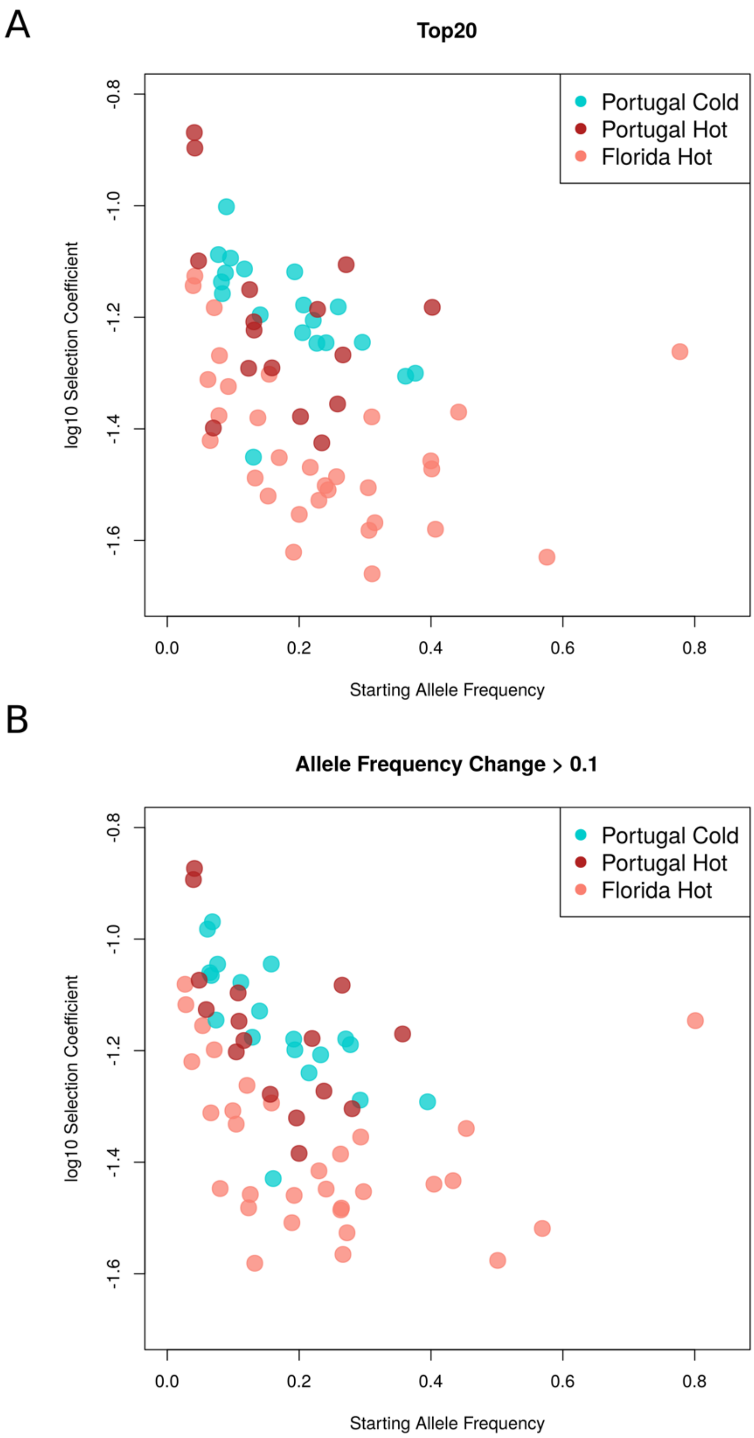


Supplementary Figure S6: The adaptive architecture is population-specific but does not depend on the temperature regime. These results were robust to the definition of the selected alleles. Selected haplotype blocks (i.e. selected alleles) from the cold-evolved Portugal (blue), hot-evolved Portugal (red) and hot-evolved Florida (pink) population are shown. A) selected alleles were defined by the top 20 % of SNPs in a selected haplotype block, B) selected alleles were defined by all SNPs with an allele frequency change > 0.1.

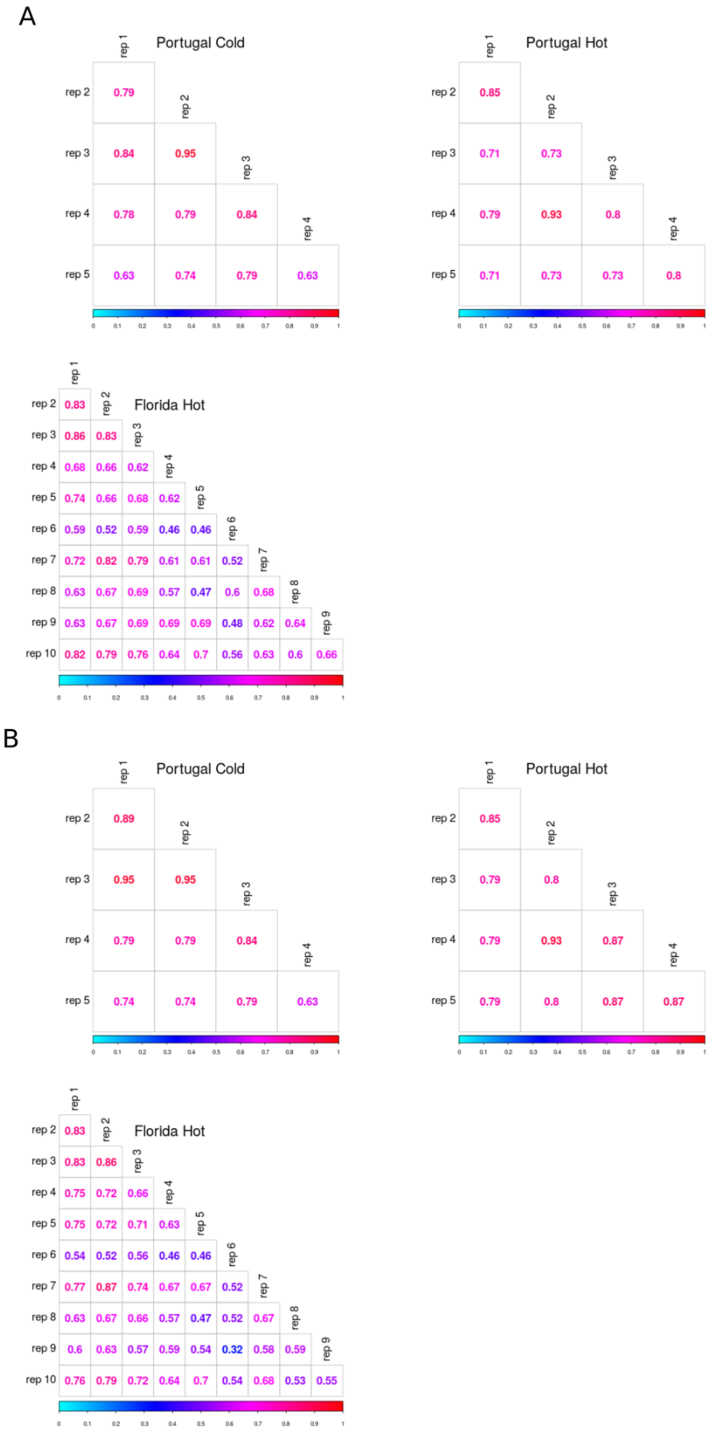


Supplementary Figure S7: Replicate populations derived from Portugal founders show a more similar selection response than those with Florida ancestry, and this observation is robust to how selected alleles were defined. Jaccard indices comparing the different replicates in the cold-evolved Portugal (top left), hot-evolved Portugal (top right) and hot-evolved Florida (bottom) population are shown. A) Jaccard index was computed based on estimation of significant selection coefficients from the top 20% SNPs (p-value < 0.05). B) Jaccard index was computed based on estimation of significant selection coefficients from the SNPs with allele frequency change > 0.1.

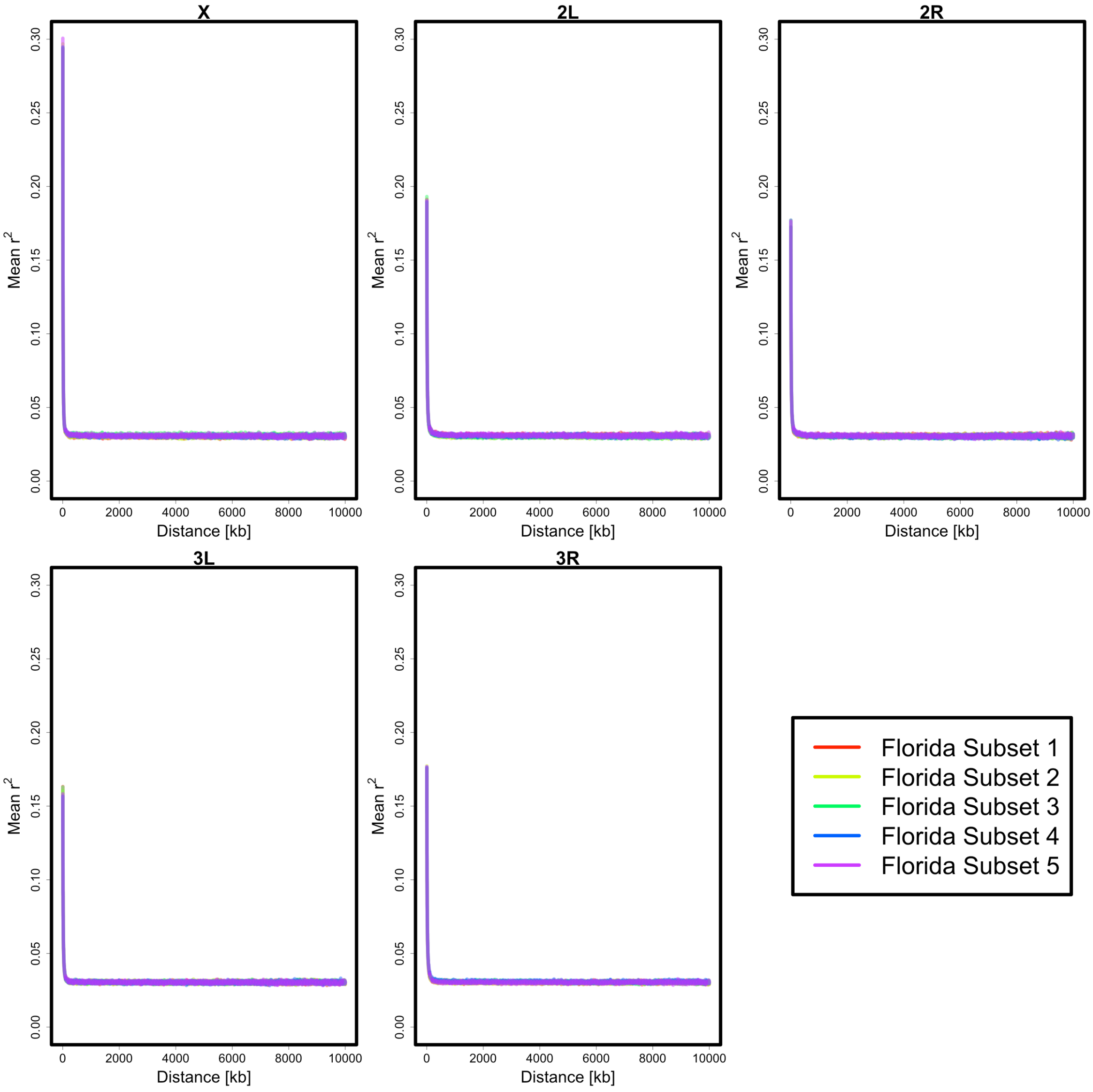


Supplementary Figure S8: Linkage disequilibrium in the ancestral Florida population as measured by the mean $r^2$ of loci with distances up to 10,000 kb is very consistent for different subsets of 34 individual haplotype sequences.
